# Presence of phage-plasmids in multiple serovars of *Salmonella enterica*

**DOI:** 10.1101/2024.02.02.574330

**Authors:** Satheesh Nair, Clare R Barker, Matthew Bird, David R Greig, Caitlin Collins, Anaïs Painset, Marie Chattaway, Derek Pickard, Lesley Larkin, Saheer Gharbia, Xavier Didelot, Paolo Ribeca

## Abstract

Evidence is accumulating in the literature that the horizontal spread of antimicrobial resistance (AMR) genes mediated by bacteriophages and bacteriophage-like plasmid (phage-plasmid) elements is much more common than previously envisioned. For instance, we recently identified and characterised a circular P1-like phage-plasmid harbouring a *bla*_CTX-M-15_ gene conferring extended-spectrum beta-lactamase (ESBL) resistance in *Salmonella enterica* serovar Typhi. As the prevalence and epidemiological relevance of such mechanisms has never been systematically assessed in Enterobacterales, in this study we carried out a follow-up retrospective analysis of UK *Salmonella* isolates previously sequenced as part of routine surveillance protocols between 2016 and 2021. Using a high-throughput bioinformatics pipeline we screened 47,784 isolates for the presence of the P1 lytic replication gene *repL*, identifying 226 positive isolates from 25 serovars and demonstrating that phage-plasmid elements are more frequent than previously thought. The affinity for phage-plasmids appears highly serovar-dependent, with several serovars being more likely hosts than others; most of the positive isolates (170/226) belonged to *S.* Typhimurium ST34 and ST19. The phage-plasmids ranged between 85.8–98.2kb in size, with an average length of 92.1kb; detailed analysis indicated a high amount of diversity in gene content and genomic architecture. 132 phage-plasmids had the p0111 plasmid replication type, and 94 the IncY type; phylogenetic analysis indicated that both horizontal and vertical gene transmission mechanisms are likely to be involved in phage-plasmid propagation. Finally, phage-plasmids were present in isolates that were resistant and non-resistant to antimicrobials. In addition to providing a first comprehensive view of the presence of phage-plasmids in *Salmonella*, our work highlights the need for a better surveillance and understanding of phage-plasmids as AMR carriers, especially through their characterisation with long-read sequencing.

**Data Summary:** All of the FASTQ files examined in this study have been uploaded to the Sequence Read Archive under BioProject PRJNA248792. Accessions of individual isolates which were found to contain phage plasmids are listed in Supplementary Table S1.

**Impact Statement:** Bacteriophage-like plasmids are increasingly being recognised as important mobile elements in many species of bacteria, particularly due to their involvement in the transmission of antimicrobial resistance (AMR); however, few studies of their overall prevalence in clinical datasets have been undertaken to date. In this study we have performed the first large-scale surveillance of human disease-associated *Salmonella* genomes for the presence of P1-like phage-plasmids, showing that they are more common than previously thought. Furthermore, we highlight how valuable information about the evolution and transmission of phage-plasmids in *Salmonella* and other Enterobacterales can be revealed by linking phage-plasmid prevalence and genetic diversity to epidemiologically relevant metadata such as *S. enterica* serovar, outbreak clusters, time, and geography. Our work shows the ability to use sequencing data and scalable bioinformatics workflows for the detection and characterisation of these extrachromosomal elements, highlights the importance of screening for novel mechanisms of AMR transmission, and provides a foundation for further surveillance studies of phage-plasmid prevalence.

## Introduction

The global threat of rising antimicrobial resistance (AMR) incidence is well known, particularly among genera of Enterobacterales such as *Escherichia* and *Salmonella* (1,2). AMR genes can be transferred between bacteria via horizontal gene transfer (HGT), which involves many diverse molecular mechanisms mediated by mobile and integrative elements including transposons and integrons, plasmids, bacteriophages and genomic islands, all of which are well known for their role in the transfer of AMR genes (3) as well as other genes such as virulence and fitness factors (4,5). Temperate bacteriophages, which typically integrate as prophages into the host chromosome, are another important mechanism of HGT and further contribute to the spread of AMR genes, toxins and virulence factors (4,6–8). Moreover, certain temperate bacteriophages have been found to exist in their prophage forms as low-copy number, extrachromosomal-plasmids and thus can replicate autonomously within bacterial cells while remaining latent (9,10).

Large-scale computational analysis of such bacteriophage-like plasmids, or “phage-plasmids”, has shown that they are associated with multiple specific types of bacteriophages including P1, P7 and SSU5 (11). The P1-like phage-plasmid community belongs to the *Myoviridae* family and forms two distinct sub-groups based on relatedness of their gene content, with one containing P1 and its relatives, and the other containing phage-plasmid D6 (10,11). The plasmid replication types linked to the P1-like phage-plasmid group are consistently either IncY or p0111 (11,12), suggesting both types can theoretically be compatible and maintained in a single host.

There is increasing evidence from the literature that the horizontal spread of AMR genes by these phage-plasmids is more common than previously envisioned, especially among the P1, SSU5 and AB types (13). It is speculated that bacteriophages carrying AMR genes would confer an evolutionary advantage to their hosts, especially during latent infection as phage-plasmids. The mechanism involved in conferring evolutionary advantage is underpinned by several complex molecular pathways currently being elucidated (14); it would eventually result in phage-plasmids with AMR genes being positively selected. For instance, phage-plasmids carrying extended-spectrum beta-lactamase (ESBL) and colistin resistance genes have been identified in *Escherichia coli* (12,15–22), *Klebsiella pneumoniae* (23) and *Salmonella enterica* (13,24–26) to date. Among *S. enterica*, reports suggest that phage-plasmids are associated more specifically with the AMR genes *bla*_CTX-M-15_ (25), *bla*_CTX-M-27_ (24,26) and *mcr-1* (26).

We recently identified, characterised and described for the first time a circular P1-like phage-plasmid harbouring a *bla*_CTX-M-15_ gene conferring high-level ESBL resistance within *S. enterica* serovar Typhi (25). Following this discovery, we wanted to determine the extent to which such elements may be distributed among all the subspecies and serovars of *Salmonella* in our collection, and whether these phage-plasmids play a significant role in the increasing spread of antibiotic resistance.

The Gastrointestinal Bacteria Reference Unit (GBRU) at the UK Health Security Agency (UKHSA) is the UK national reference laboratory for enteric bacterial pathogens. Routine whole genome sequencing (WGS) for all *Salmonella* spp. was implemented in 2014, resulting in a collection of roughly 9000-10000 genomes per year from hundreds of different serovars. In this study, we offer a retrospective examination of the genomic data associated with this national surveillance. We developed a high-throughput process for detecting the occurrence of P1-like phage-plasmids and applied it to examine the genomes of *Salmonella* spp. isolates received by the reference laboratory. Our goal was to gain insight into the distribution, prevalence, and connection of these phage-plasmids with antimicrobial resistance genes.

## Materials and Methods

### Data collection & bacterial strains

A total of 47,784 *Salmonella* spp. isolates were sequenced by GBRU between January 2016 and December 2021. All of the FASTQ files have been uploaded to the Sequence Read Archive under BioProject PRJNA248792, where links to automatically generated genome assemblies and annotated nucleotide sequences can also be found (Table S1). Ethical approval for the detection of gastrointestinal bacterial pathogens from faecal specimens, or the identification, characterisation and typing of cultures of gastrointestinal pathogens, submitted to GBRU is not required as it is covered by the surveillance mandate of UKHSA.

### Whole genome sequencing and bioinformatics

Genomic DNA was extracted using the Qiasymphony DSP DNA Midi Kit on the Qiasymphony system (Qiagen, UK) to manufacturer’s instructions. The Nextera XP kit (Illumina) was used to prepare the sequencing library for sequencing on the HiSeq 2500 and NextSeq 1000 instruments (Illumina), run with the fast protocol. Trimmomatic v0.27 (27) was utilised to remove bases with a PHRED score of <30 from the leading and trailing ends on the FASTQ reads, with reads <50bp after trimming discarded. The eBurst Group (eBG) was determined as previously described (28). Multilocus sequence type (MLST) assignment was performed using MOST v2.18 (29), and the FASTQ reads underwent single nucleotide polymorphism (SNP) typing and clustering using SnapperDB v0.2.9 (30), which additionally produced SNP addresses for the isolates: these represent the relationship between isolates using hierarchical single linkage clustering of pairwise SNP distances at several thresholds (250, 100, 50, 25, 10, 5 and 0 SNPs, respectively). Routine detection of AMR genes and plasmid replication gene types was conducted using in-house GastroResistanceFinder v2.7, which derives from GeneFinder (available at https://github.com/ukhsa-collaboration/gene_finder).

### Detection of *repL* gene

The *repL* gene forms part of the L-replicon of the P1 family of temperate bacteriophages, which is the active replicon for DNA replication during the lytic cycle (9); therefore, we used the presence of *repL* as an indicator for carriage of a P1-like phage plasmid. For each of the 47,784 samples, reads were mapped to the *repL* gene sequence with the GEM mapper v3.6.0 (31) in default mode. Alignments were filtered to keep those of sufficient quality (having no more than 2 indels or clippings, clippings being defined as CIGAR operation “S” in the SAM format (32) documentation – available at https://github.com/samtools/hts-specs – such that the sum of the number of substitutions, indels and clippings would not exceed 5% of the read length; and such that the sum of the number of mismatches and the lengths of indels and clippings would not exceed 10% of the read length); all multimaps were kept. Pileups were generated with Samtools (32,33) pileup v1.15.1 (using htslib (34) v1.16) and parsed to determine coverage. The distribution of the fraction of gene bases having non-zero coverage was observed to be bimodal, with peaks at either very small fractions (likely corresponding to sequencing noise) or very high fractions (likely corresponding to true positives). We considered a sample to be a potential positive for the detection of *repL* whenever the number of gene bases supported by sequencing reads was >60% of the total, which coincides with the position of the valley between the two peaks of the distribution. While we did not observe correlation between the fraction of gene bases supported by sequencing reads and read coverage (i.e., there were phage-plasmids with very low read coverage but a complete *repL* gene) we further required the average coverage of bases having non-zero coverage to be >1, to ensure that on average each gene base was supported by more than one single sequencing read. This procedure produced 248 candidates. Further manual inspection based on metadata yielded 226 candidate samples (Table S1).

### Genome assembly and phylogenies

The 226 genomes in which the *repL* gene was present with high confidence according to the alignment pipeline were assembled *de novo* from the paired-end reads using SPAdes v3.8.0 (35). The assembled genomes were visualised using Bandage v0.8.1 (36), with the built-in BLAST (37) tool used to identify the phage-plasmid contig(s) associated with the *repL* gene (NC_005856.1) and the IncY *repA* (K02380.1) and p0111 *repB* (AP010962.1) plasmid replication genes, as well as to determine the location of any AMR genes (using the Resfinder v4 database (38)). The putative phage-plasmid contigs were manually curated, extracted and used to perform all downstream comparative genomic and phylogenetic analyses. A total of 33/226 genome assemblies were of too poor quality to extract the phage-plasmid contigs for further analysis, as determined by manual inspection in Bandage and BLASTn against the P1 reference (NC_005856.1) revealing them to be spread across >5 contigs; we were unable to accurately determine their full sequence or to extract core gene content. Panaroo v1.3.4 (39) was used to produce core gene alignments (present in >98% of the remaining 193 phage-plasmids, as well as selected publicly available phage-plasmid sequences (Table S2)), which were passed to FastTree v2.1.11 (40) to generate maximum-likelihood trees using the GTR CAT method of nucleotide substitution and 100 bootstrap replicates. Trees were visualised alongside the metadata using Microreact (41).

Of the 193 phage-plasmid sequences, 39 were determined using Bandage to have assembled into single, circular contigs, while a further 69 were single, linear contigs. For individual analyses, phage-plasmid contigs were annotated using Bakta v1.8.2 (42), and visualised and compared using Easyfig (43) and Clinker (44). To examine the taxonomic context of the phage-plasmids we compared a representative subset of 25 sequences from a range of serovars to the full collection of prokaryotic dsDNA viruses on the ViPTree server (45), producing a proteomic dendrogram based on tBLASTx genome-wide sequence similarity.

### Phylogenetic analysis of plasmid replication type

To examine how the plasmid replication type varied across the *repL*-positive phage-plasmid tree, we performed ancestral state reconstruction (46) and a phylogenetic randomisation test. First, to determine if the two replication types were distributed randomly across the tips of the tree without regard to tree structure, we performed a simple label-switching test. The observed states, IncY (n=79/193) and p0111 (n=114/193), were randomly reassigned to new terminal nodes. This process was repeated 100 times. In each case, a Welch t-test was performed using the t.test function in R package *stats* v 4.3.1 (47) to assess whether the distribution of Euclidean genetic distances between IncY and p0111 individuals differed between the observed and the tip-randomised dataset. The mean Bonferroni-corrected p-value is 1.07e-15, indicating that plasmid replication type is not distributed randomly across the tips of the tree if one disregards tree structure.

However, as the presence of large clades with near-zero branch lengths might inflate the appearance of clustering and vertical inheritance in the replication type, we needed to account for phylogenetic structure. We performed ancestral state reconstruction using the discrete maximum-likelihood method via the functions pml and ancestral.pml in R package *phangorn* v 2.11.1 (48) to infer the likely replication type state on the internal branches of the phage-plasmid tree, identifying 17 changes in replication type along the tree (Fig. S1). To determine whether these 17 changes occurred randomly across the phylogenetic history of the sample, we performed a phylogenetically-informed randomisation test. The 17 changes were randomly reassigned to new branches, with probability of reassignment proportional to branch length, and IncY/p0111 states were simulated from root to tips according to these change-points with the phen.sim function in R package *treeWAS* v 1.1 (49). We performed 100 simulations and repeated the Welch t-test as described above to determine whether the observed replication type states showed any greater propensity for vertical inheritance than one might expect despite the number of changes inferred.

## Results

### Phage-plasmid prevalence

Using our mapping pipeline, the *repL* gene was detected in the reads of 226 isolates (0.47%), comprising 0.2% to 1.1% of the dataset depending on the year (Table 1, Table S1). The *repL* gene was present in 25 different *Salmonella* serovars and was most prevalent among isolates of *S.* Typhimurium (n=170), followed by *S.* Oslo (n=9), *S.* Livingstone (n=6), and *S.* Indiana (n=5). Table 1 also shows serovar prevalence for the whole set of 47,784 isolates. Comparison of serovars for which phage-plasmids were detected suggests that they tend to occur much less than expected in *S.* Enteritidis (0.44% with phage-plasmids vs. 27% prevalence of *S.* Enteritidis within our whole *Salmonella* collection); less than expected in *S.* Agona (0.9% vs. 2.3%), *S.* Infantis (1.3% vs. 4.3%), *S.* Typhi (0.9% vs. 3.2%); much more than expected (84% vs. 19%) in *S.* Typhimurium, *S.* Oslo (4.0% vs. 0.31%), and *S.* Livingstone (2.7% vs. 0.21%); and more than expected in *S.* Indiana (2.2% vs 0.9%), *S.* Mbandaka (1.8% vs. 0.81%), and *S.* Mikawasima (1.8% vs. 0.76%). Phage-plasmids were also not detected in a large number of serovars often occurring in the full surveillance datasets (the most frequent being *S.* Newport, corresponding to 3.3% of the 47,784 isolates; *S.* Paratyphi A, 1.6%; *S.* Kentucky, 1.5%; and *S.* Braenderup, 1.5%).

**Table 1–.** Summary of metadata for the 226 *Salmonella* isolates possessing the *repL* gene; see Table S1 for individual isolate information. Values within brackets indicate frequency of the Rep type(s). In the column “Prev. surv.” (“prevalence in surveillance”) we report the percentage of occurrences in the 47,784 *Salmonella* spp. isolates that were sequenced by GBRU between January 2016 and December 2021; in the column “Prev. phm.” (short for “prevalence in phage-plasmids”) we report the percentage of occurrences in the set of sequences that were found to contain phage-plasmids as a result of the current study.

| Serovar | No. | Prev. phm. | Prev. surv. | ST(s) | Years | Rep type(s) | AMR |
| --- | --- | --- | --- | --- | --- | --- | --- |
| Typhimurium | 170 | 84% | 19% | 34, 19 | 2015-2021 | IncY (63)<br>p0111 (107) | None |
| Oslo | 9 | 4.0% | 0.31% | 1370 | 2016 | IncY (5)<br>p0111 (4) | None |
| Livingstone | 6 | 2.7% | 0.21% | 457, 638 | 2018-2019, 2021 | IncY (4)<br>p0111 (2) | None |
| Indiana | 5 | 2.2% | 0.90% | 17 | 2018 | p0111 | None |
| Mbandaka | 4 | 1.8% | 0.81% | 413, 1602 | 2016, 2018, 2020 | IncY (3)<br>p0111 (1) | None |
| Mikawasima | 4 | 1.8% | 0.76% | 1815 | 2016-2018 | IncY | None |
| Virchow | 4 | 1.8% | 1.4% | 16 | 2016-2018, 2020 | IncY (2)<br>p0111 (2) | None |
| Infantis | 3 | 1.3% | 4.3% | 32 | 2018 | p0111 (3) | None |
| Stanley | 3 | 1.3% | 1.6% | 29 | 2018-2019 | IncY (3) | None |
| Agona | 2 | 0.90% | 2.3% | 13 | 2019 | p0111 (2) | None |
| Typhi | 2 | 0.90% | 3.2% | 1 | 2019 | IncY (2) | *bla*<sub>CTX-M-15</sub> |
| Adjame | 1 | 0.44% | 0.11% | 3929 | 2021 | IncY | None |
| Bareilley | 1 | 0.44% | 1.2% | 203 | 2018 | p0111 | None |
| Corvallis | 1 | 0.44% | 0.42% | 1541 | 2016 | p0111 | None |
| *S. enterica* 45:b:- | 1 | 0.44% | 0.66% | 2524 | 2016 | IncY | None |
| Enteritidis | 1 | 0.44% | 27% | 11 | 2016 | IncY | None |
| Florian | 1 | 0.44% | 0.002% | 7166 | 2020 | p0111 | None |
| Florida | 1 | 0.44% | 0.021% | 351 | 2017 | IncY | None |
| Minnesota | 1 | 0.44% | 0.31% | 548 | 2019 | IncY | None |
| Mississippi | 1 | 0.44% | 0.25% | 4711 | 2020 | IncY | None |
| Monschau | 1 | 0.44% | 0.08% | 443 | 2017 | IncY | None |
| Ohio | 1 | 0.44% | 0.12% | 329 | 2017 | IncY | None |
| Pomona | 1 | 0.44% | 0.084% | 451 | 2016 | p0111 | None |
| Rissen | 1 | 0.44% | 0.19% | 469 | 2017 | IncY | None |
| Telaviv | 1 | 0.44% | 0.0063% | 1068 | 2016 | p0111 | None |

### Phage-plasmid features

To explore the structure and content of the putative phage-plasmids, we performed *de novo* assembly of the 226 *repL*-positive *Salmonella* genomes and extracted the contigs upon which the *repL* gene was located, where possible (n=193). Analysis of contigs assessed to be circular (n=39) showed that the phage-plasmids ranged between 89.8–97.6kb in size with an average length of 93.1kb. Expanding this to include all phage-plasmids that had assembled into a single circular or linear contig (n=108) gave a range of 85.8-98.2kb (average 92.1kb), except for four isolates of *S.* Typhimurium which had notably shorter or longer phage-plasmid sequences of 76-79kb and 104-109kb (Table 1, Table S1). Additionally, two other *S*. Typhimurium genomes had phage-plasmids located on single contigs >115kb in length; these were putatively determined to instead be integrated into chromosomal sequences of these isolates (Fig. S2); long-read sequencing would be necessary to corroborate this finding.

A search against a database of plasmid replication genes revealed that 94 of the phage-plasmids had the IncY plasmid replication type and 132 had the p0111 replication type. To assess the level of vertical or horizontal transmission from the correlation between phage-plasmid phylogeny and plasmid replication type, a maximum-likelihood tree based on a 51,260bp alignment of the 54 core genes from the 193 phage-plasmid contigs was constructed (Fig. 1). Statistical analysis showed that the two types are not distributed completely randomly across the tips of the tree, if one disregards the tree structure (p-value = 1.07e-15). That excludes a scenario where transmission occurs in a fully horizontal manner. We then conducted a different analysis that considers the structure of the phylogenetic tree. Ancestral state reconstruction (46) revealed that 17 changes in replication type had occurred along the evolutionary history of the phage-plasmid sample (Fig. S1). Comparison to simulated data suggested that these changes occur randomly, falling with greater probability on longer branches, and result in a distribution of replication types that does not differ statistically from a random distribution once branch lengths are considered (p-value = 0.15). Vertical inheritance of replication type thus occurs among clones and some close relatives, but it is not maintained over greater genetic distances. This supports a mixed model of transmission, whereby, as expected, both vertical and horizontal gene transfer mechanisms play a role.

**Fig. 1–.**
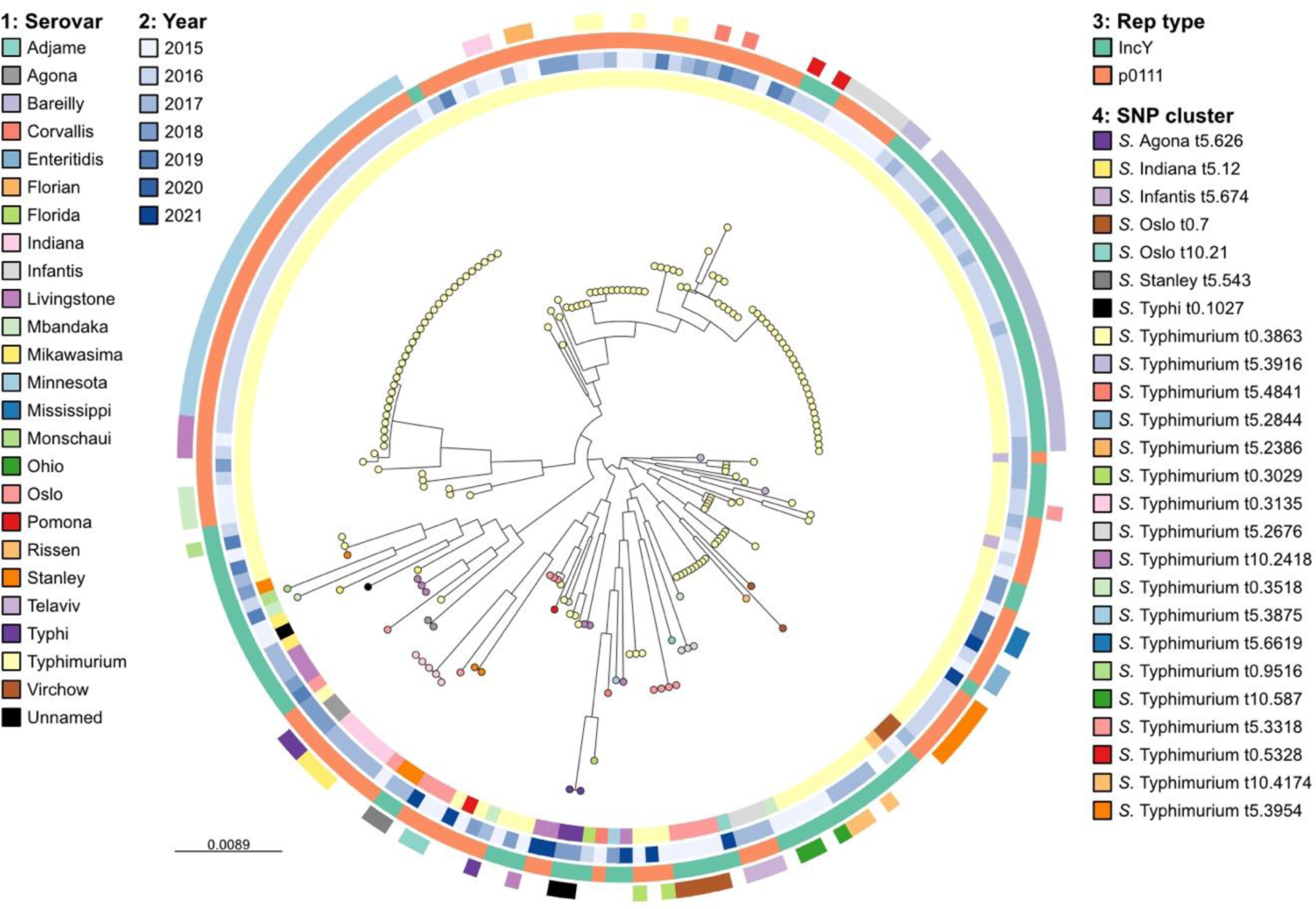
Midpoint-rooted maximum-likelihood tree of 51,260bp alignment of 54 core genes from the 193 *repL*-positive phage-plasmid contigs, with tips coloured by serovar. Rings from inner to outer: serovar; year; replication gene type; host isolate SNP cluster based on 0-10 SNP threshold of SNP address (Table S1).

### Genetic diversity

In almost all cases, phage-plasmid sequences are grouped according to the host isolate SNP clustering (as determined by 0-10 SNP thresholds, described by the SNP addresses in Table S1) but are not necessarily closely related to others from the same serovar (Fig. 1, Fig. 2). However, there are examples of cross-serovar similarity, with a phage-plasmid from an *S.* Stanley that is highly similar to several from *S.* Typhimurium (Fig. S3). Additionally, there are also phage-plasmids from closely related, epidemiologically-linked *S.* Typhimurium strains (forming a 10-SNP cluster based on SNP address) that display considerable genetic diversity and even possess different plasmid replicon types (Fig. 3). In relation to the P1 phage-plasmid itself, several regions are variable (Fig. S4) including: 47.5kbp - 53.6kbp (encoding a type 3 restriction-modification enzyme); 65.7kbp - 68.1kbp (encoding autolytic and phage proteins); 76.3kbp - 78.9kbp (encoding phage tail fibre proteins). However, there are large sections of conserved sequence surrounding the previously mentioned variable regions which contain a variety of phage and plasmid associated genes.

**Fig. 2–.**
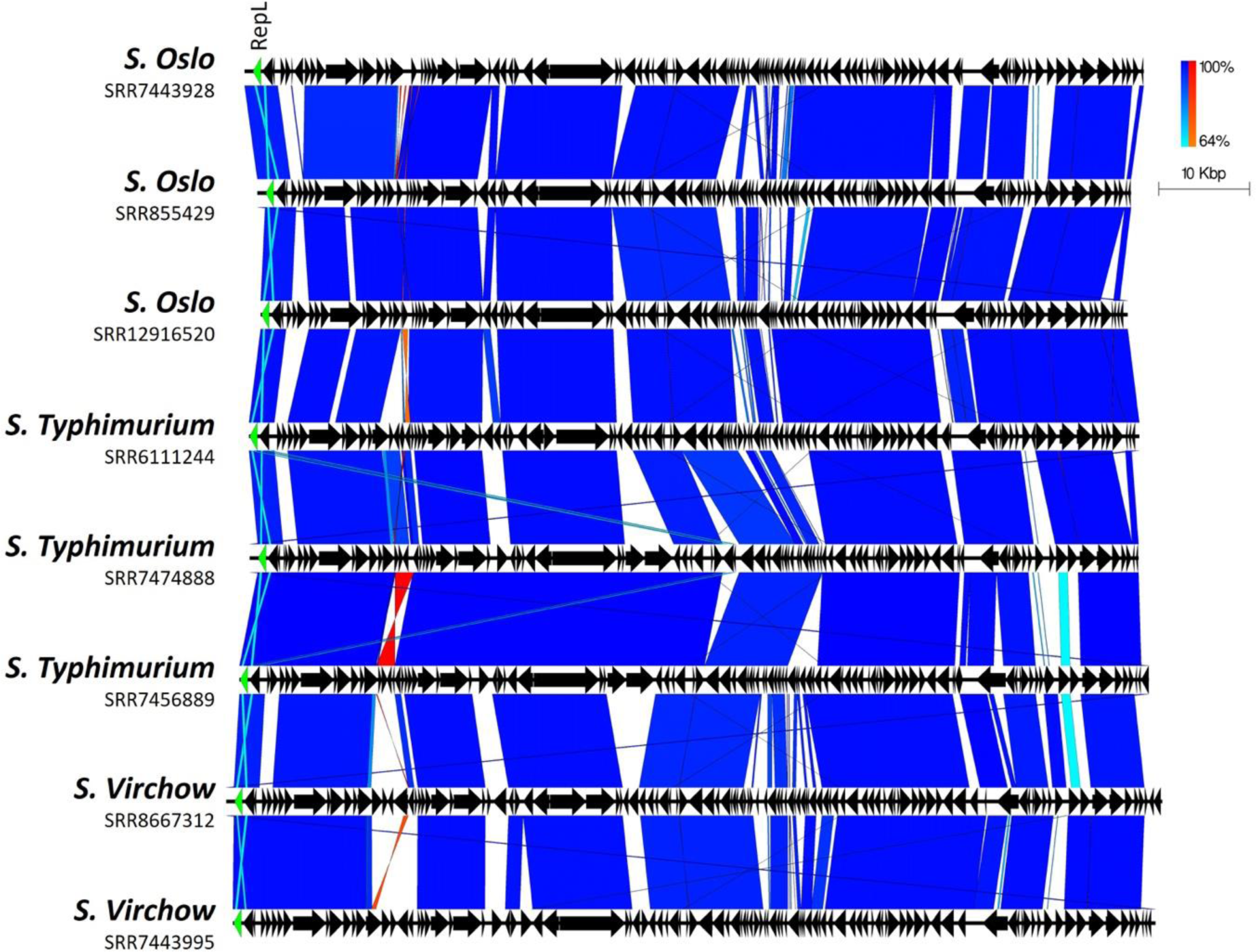
Plot showing how diversity occurs in sections in-between highly conserved regions, even among isolates belonging to the same serovar. Arrows indicate gene direction; scale bars indicate level of nucleotide similarity for forward (blue) and reverse (red) sequences. The *repL* gene is highlighted in green.

**Fig. 3–.**
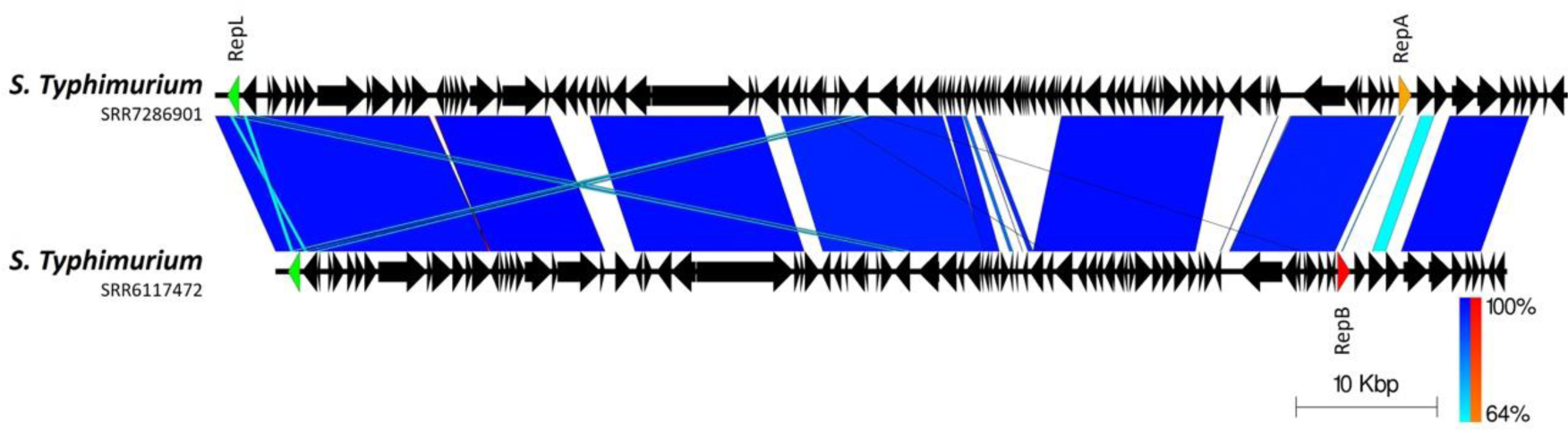
Plot showing variability in the sequences of two phage-plasmids from *S*. Typhimurium isolates that belong to the same 10-SNP cluster. Arrows indicate gene direction; scale bars indicate level of nucleotide similarity for forward (blue) and reverse (red) sequences. The *repL* gene is highlighted in green; the IncY *repA* in orange and the p0111 *repB* in red.

We next performed a BLASTN search of the GenBank database to identify other phage-plasmids that were related to those in our collection, including from other species of Enterobacterales. We produced a tree containing selected representative phage-plasmid sequences from our collection, alongside 31 publicly available sequences labelled as either phage-plasmids, plasmids or bacteriophages (Table S2). This confirmed the high amount of genetic diversity and additionally showed that there is no apparent correlation between phage-plasmid sequence and species (Fig. 4). We did however identify several sequences (typically labelled as either plasmid or bacteriophage) that are highly similar to phage-plasmids from our collection (Fig. S5). To further examine the taxonomic context of our phage-plasmid dataset, we compared 25 representative sequences taken from a range of serovars to a large public collection of prokaryotic dsDNA viruses, confirming that the P1-like phage-plasmids are monophyletic, and that the closest relative to this clade is *Escherichia* phage D6 (inset Fig. 4, Fig. S6).

**Fig. 4–.**
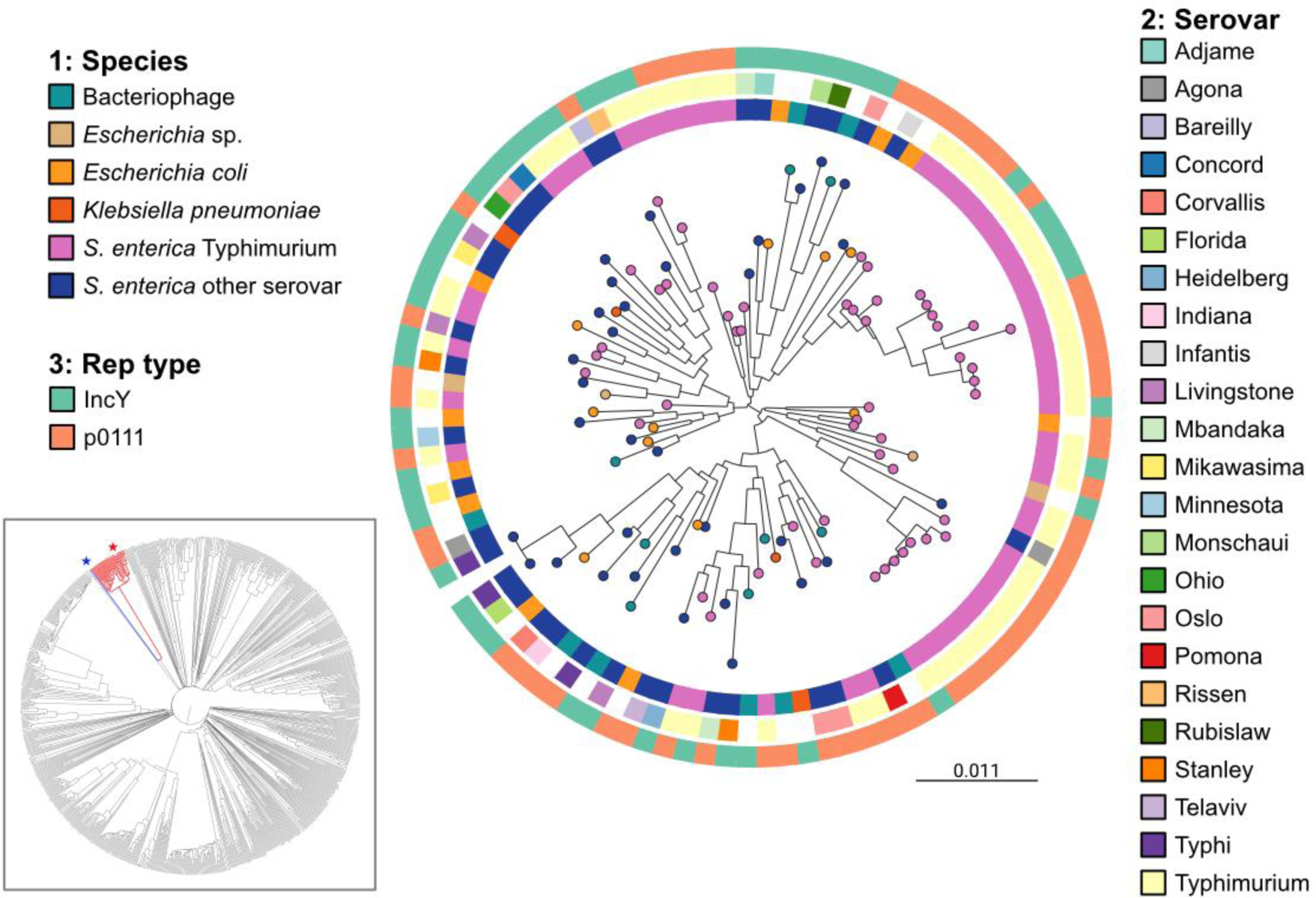
Midpoint-rooted maximum-likelihood tree of 42,606bp alignment of 46 core genes from 107 phage-plasmids, including 31 from public databases, with tips coloured by species. Rings from inner to outer: species; *S. enterica* serovar; replication gene type. Inset figure: unrooted proteomic tree of 825 prokaryotic dsDNA viruses from the ViPTree server (45) showing the single clade containing P1-like phage-plasmids (including P1, P7, SJ46 and 25 from this study) coloured in red. Phage P1 is highlighted with a red star, and phage D6 by a blue star.

### Antimicrobial resistance

AMR genes were detected in all 170 of the *S.* Typhimurium isolates and in 25/57 of the isolates belonging to other serovars (Table S1). Preliminary visual assessment of the whole genome sequences indicated that the majority of AMR genes were present on the chromosomes of these isolates rather than being associated with the phage-plasmid contigs. However, we did confirm the presence of *bla*_CTX-M-15_ genes encoding ESBL resistance on phage-plasmids from two *S.* Typhi isolates (Fig. 5), both of which were associated with travel to Iraq. We have previously reported and described the phage-plasmid sequence in detail for one of these isolates (25). Further comparison with publicly-available plasmid and phage-plasmid sequences showed that the flanking region of *bla*_CTX-M-15_ is variable, and that the gene is located next to an IS*Ec9*/IS*Ecp1* element in the *S.* Typhi isolates that is similar to that alongside the *bla*_CTX-M-55_ gene of *E. coli* phage-plasmid JL22 (ON018986), but different to that next to the *bla*_CTX-M-27_ in *S*. Indiana phage SJ46 (KU760857).

**Fig. 5–.**
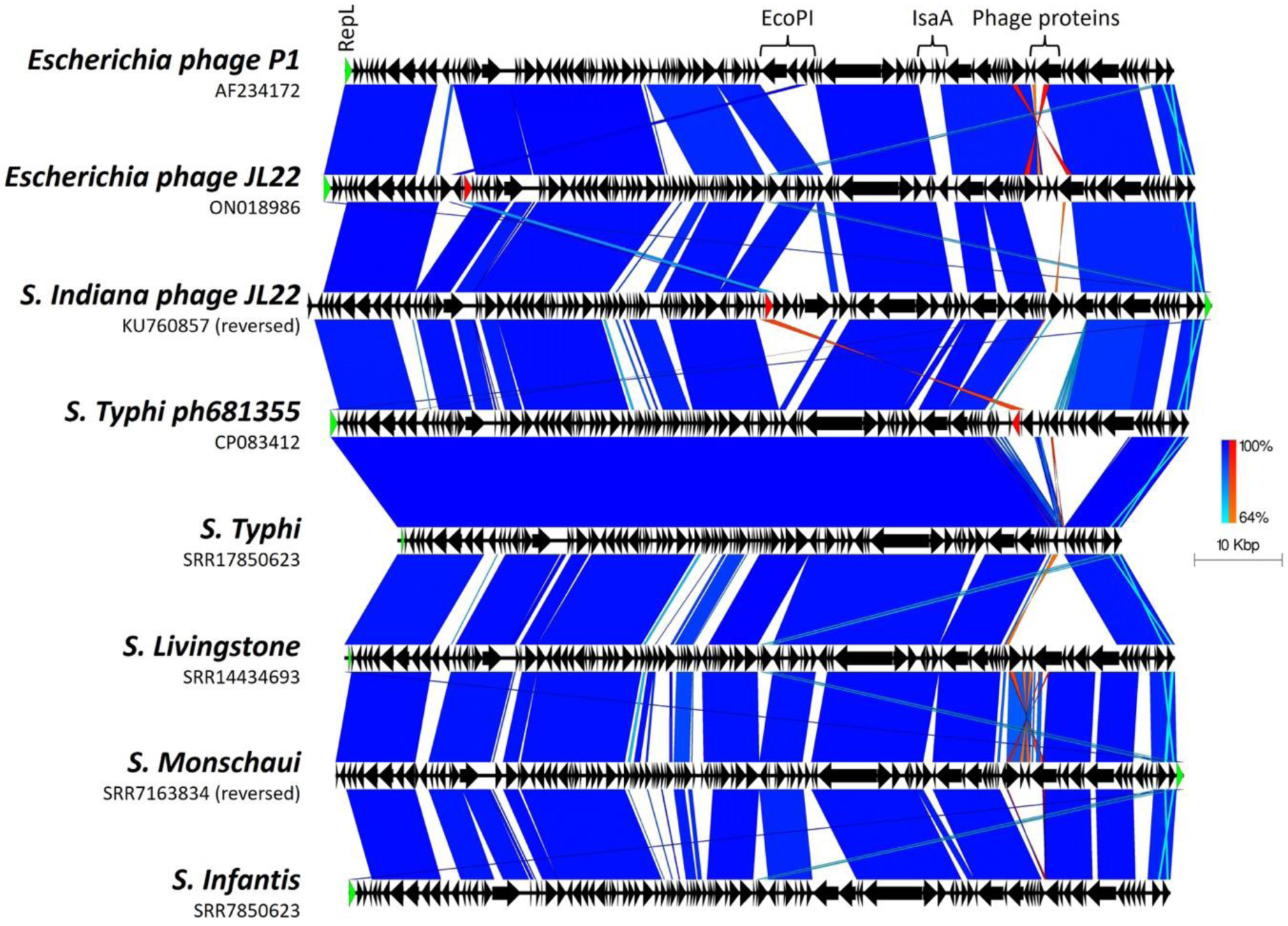
Comparison of *S*. Typhi ESBL-carrying phage-plasmid (CP083412: *bla*_CTX-M-15_) to other ESBL-carrying phage-plasmids (ON018986: *bla*_CTX-M-55_ & KU760857: *bla*_CTX-M-27_). Arrows indicate gene direction, scale bars indicate level of nucleotide similarity for forward (blue) and reverse (red) sequences. The *repL* gene is highlighted in green. In the figure we show that those encoding ESBL genes (CP083412, ON018986 & KU760857) shown by the red arrows, are encoded within variable loci, and not within the same region.

## Discussion

New and emerging research based on WGS highlights the prevalence of many different phage-plasmids in the bacterial world (11,12,19,26). We performed retrospective genomic surveillance for the presence of P1-like phage-plasmids within a collection of almost 48,000 *Salmonella* isolates received over a five year period. Our work shows that phage-plasmids are distributed among multiple serovars of *Salmonella* and have been present within the UK population since at least 2016. While there is growing evidence of a significant phage-plasmid presence among genera of Enterobacterales (11,13,19), targeted large-scale genomic surveillance of *Salmonella* had not previously been performed.

P1-like phage-plasmids have been reported in a small number of individual strains of *Salmonella* (24–26), however previous large-scale genomic surveillance studies of the Enterobacterales have primarily focused on *E. coli*. Our work shows that the overall prevalence of P1-like phage-plasmids among the clinical serovars of *S. enterica* appears to be considerably lower than in *E. coli*, being just ∼0.5% compared to 7-12.6% (15,19). However, this may vary depending on country or isolate source, as a higher prevalence has been found in *Salmonella* from pork in China (24). Interestingly, this study also suggested an association between the phage-plasmid sequences and their host strains, with closely related *S. enterica* isolates (<5 SNPs) possessing identical or extremely similar phage-plasmids; though only among specific strains of *S.* Derby and *S.* Indiana that were sampled from the same facility. The difference in prevalence between our collection and studies such as theirs may be a result of the clinical bias that is inherent in our dataset. We saw hints of this when selecting public phage-plasmid sequences for comparison: many of the P1-like sequences from *S. enterica* and other Enterobacterales that are available online were sampled from animals, food, or the environment. It would therefore be beneficial to extend such screening to other large genomic (and metagenomic) datasets that contain comparatively more food and environmental samples to gain an accurate view of overall prevalence in *S. enterica*.

We confirmed the presence of P1 phage-plasmids in multiple *Salmonella* serovars, with the majority (84%) occurring in *S.* Typhimurium. This represents a significantly higher fraction than the fraction of *S.* Typhimurium recorded in our surveillance database during the same time period. Interestingly, it has been reported that isolates belonging to *S.* Typhimurium have a higher prophage burden than many other serovars of *Salmonella* (50), possibly because phage-mediated recombination plays a role in the periodic emergence of epidemic clones of *S.* Typhimurium (8). We also found in this study that other serovars seem to be especially amenable to acquiring other such elements – examples are *S.* Oslo, *S.* Livingstone and *S.* Indiana. Perhaps even more importantly, we notice that some serovars very rarely acquire phage-plasmids, with the fraction of isolates containing phage-plasmids being much smaller than what would be expected from the fraction of surveillance isolates. That is especially true for *S.* Enteritidis (0.44% vs. 27%, or ∼50 times less); other examples are *S.* Agona, *S.* Infantis and *S.* Typhi. While the exact mechanism of affinity between serovars and phage-plasmids is likely to depend on several factors, we speculate that a broad link with the level of host specificity of an organism might exist, with generalists like *S.* Typhimurium and *E. coli* requiring a higher level of genome adaptability via acquisition of malleable extrachromosomal mobile elements; though this does not explain the low prevalence observed in *S.* Enteriditis.

Similar to previous studies of P1-like phage-plasmids, we report that the majority of those detected in our dataset fall into a narrow size range of 85.8-98.2kb, with the 92.1 kb average matching well to those from other collections. Others have suggested that this small size range may be due to a limit on the amount of genetic material that can be packaged within the virion head, although there have been reports of larger phage-plasmids ranging up to 155kb (51). The structure of the P1 phage-plasmids when comparing between *Salmonella* strains are diverse, but nonetheless share a common genomic architecture. Our results, alongside those of other studies (12,24), highlight large regions of conserved sequence interspaced with more variable regions, which mainly encode hypothetical genes and phage related genes rather than plasmid coding sequences. However, the conserved regions also contain phage genes along with some plasmid associated genes.

Based on SNP typing of the corresponding *Salmonella* chromosomes there is an association between the phage-plasmid sequences and chromosomal diversity, with closely related (<10 SNPs) isolates possessing identical or highly similar phage-plasmids. We also detected several P1-like phage-plasmids in online databases that share considerable sequence similarity with specific sequences from our collection. These public phage-plasmid sequences derive from various sources and span multiple years, continents and even host species; implying widespread horizontal transmission. Another study also detected an identical phage-plasmid that was present in two serovars of *S. enterica* sampled from the same facility (24). In combination with the link to chromosomal relatedness this suggests that horizontal transfer may be followed by maintenance of the phage-plasmids during clonal expansion of strains, with subsequent loss of the phage-plasmid occurring sporadically in some isolates, rather than there having been long-term vertical transmission and evolution at a serovar or species level. This is reinforced by our discovery that there is only limited phylogenetic clustering of the two different plasmid replicon types: one type is maintained across each shallow clade but frequent switching in replication type breaks this association when considering deeper lineages. Additionally, our results suggest that overall genetic diversity of the phage-plasmids is not consistent with the corresponding population structure of their hosts at either a serovar or species level, which has also been observed in other studies (52) and is indicative of their extensive recombination and horizontal method of acquisition.

Phage-plasmids are known to be involved in the horizontal transmission of AMR genes, antiseptic resistance genes (13) and virulence factors such as toxins (4), as well as being involved in bacterial survival (51). The P1-like family in particular is the most common type of phage-plasmid to be associated with AMR gene presence (13), with a prevalence that lies between that of phages and plasmids. In our collection, P1-like phage-plasmids were present both in isolates that were genotypically resistant and susceptible to antibiotics, with patterns depending on the serovar. Most of these AMR genes were observed to be located on the chromosome, and so while it is difficult to be certain without utilising long-read sequencing, we did not find an association with AMR in almost all of the phage-plasmids in this study. However, we previously described an IncY P1-like phage-plasmid carrying *bla*_CTX-M-15_ (25) and in our current study we identified an identical phage-plasmid sequence, also possessing the *bla*_CTX-M-15_ gene, in an isolate that is genetically closely related (0 SNPs distance) as well as epidemiologically connected (same travel location). This finding therefore provides further evidence of either transduction of the AMR-carrying phage-plasmid between these two isolates, or of maintenance of this phage-plasmid during clonal expansion (while likely under the same selective pressures). Importantly, our previously described antimicrobial susceptibility testing indicated phenotypic ESBL resistance, confirming the functionality of the AMR genes present on phage-plasmids. The flanking regions and associated IS elements responsible for mobilisation of *bla*_CTX-M_ gene variants are widely discussed in the literature (53,54). The two *S.* Typhi phage-plasmids described here possess *bla*_CTX-M-15_ flanked by IS*Ec9*/IS*Ecp1*, which are also implicated in many plasmids carrying various ESBL resistance genes from multiple species (55,56); further evidence that phage-plasmids can horizontally acquire AMR genes from diverse sources (52).

The presence of AMR-conferring genes in phage-plasmids lends itself to a straightforward evolutionary explanation, whereby latent phage-plasmids would increase the fitness of their host when antibiotics are present in the environment. However, many of the phage-plasmids we identified lack genes with an explicit or known AMR function, and mostly contain unannotated genes. The accessory content of temperate phages within *Salmonella* and other bacterial genera is increasingly being revealed to provide important effects for phages as well as their hosts; such as self-defence systems that protect against infection by other phages (57,58), increased pathogenicity (59–62), and improved fitness, biofilm formation and stress responses (59,63,64). Active surveillance of their appearance and distribution might be a valuable tool to understand how bacteria harbouring phage-plasmids will evolve in the future. The unknown genes carried by phage-plasmids might act as a reservoir of foreign material, allowing a rapid response to selective environmental pressures such as antimicrobials, host defence mechanisms or available substrates. Our previously described spontaneous loss of the *bla*_CTX-M-15_ module from the *S.* Typhi phage-plasmid (25) may be evidence of such a dynamic nature. It is also possible that P1-like phage-plasmids are able to be maintained at a relatively low fitness cost to the bacterium due to their low copy number and restricted size. Furthermore, as prophages have been implicated in driving the evolution of *Salmonella* via re-assortment of virulence and fitness factors to form new pathogenic variants (65), it is possible that phage-plasmids too may play a role in this process. Other recent work has also interestingly highlighted phage-plasmids – in particular P1-like phage-plasmids – as mediating gene flow between phages and plasmids, resulting in the creation of novel elements (48).

Our results would not be possible without the creation of a robust and scalable workflow that allowed us to reprocess in one hour short-read sequencing data from hundreds of isolates on a single high performance computing (HPC) node. Our method allows an arbitrary number of genes to be detected, with high sensitivity and accuracy, and, given a sufficient computational budget, it can be scaled up to an essentially arbitrary number of samples. The *repL* gene has previously been used as a target for screening using both *in vitro* (12,24,26) and *in silico* (19,26) based methods. We build on these to prove that *repL* presence is a valid detection method to quickly screen very large genomic datasets for the presence of P1-like phage-plasmids. A previous study did however detect an atypical P1-like phage-plasmid that was missing its *repL* gene (19), suggesting that additionally targeting the IncY/p0111 replication genes may enhance the robustness of the method.

The P1-like family of phage-plasmids was the target of this work due to its proven links with AMR gene carriage, though we are determined to expand the scope of our surveillance to include screening other phage-plasmid types including SSU5 (66,67) that are known to encode AMR genes (13,68), as well as other enteric pathogens received by the UK reference laboratory such as *E. coli* and *Shigella* spp.. Active surveillance of plasmids, plasmid replicon types and AMR gene presence is already part of our routine bioinformatics pipeline, and so the implementation of screening for phage-plasmids would be a valuable complement to this service. Additionally, this method could be used to rapidly screen publicly available genome collections, for example on the Sequence Read Archive (69) or Enterobase (70), to get a more accurate idea of global P1-like phage-plasmid prevalence within *Salmonella* and other species. Finally, future studies of phage-plasmids will benefit from the application of techniques, such as long-read sequencing, that are able to capture much more easily the structure and genomic architecture of complex plasmid-like elements, including those carrying AMR modules.

## Conclusions

We developed and applied a robust, high-throughput method to perform the first large-scale genomic surveillance of P1-like phage-plasmids in a national collection of *Salmonella* isolates. Although the prevalence is lower than what has been reported for other species of Enterobacterales, the phage-plasmids we identified are present across a number of serovars. They carry genes conferring not only AMR, but also several so far unknown and potentially relevant traits. The apparent ease of transmission across serovars, species and continents, and potential to integrate into the chromosome, highlight the importance to routinely monitor the presence of such phage-plasmids. We have expanded the knowledge of P1-like phage-plasmids by linking their sequences back to host genomic diversity and supplementing them with epidemiological metadata, showing that while such phage-plasmids are understood to be ancient features, their acquisition by bacterial strains appears to be a dynamic process driven by recent evolutionary pressure and so it is important to examine them in the context of their host strains. This study highlights the necessity of regular sequence data curation for the detection, characterisation and tracking of novel mobile elements involved in AMR transmission and other factors that can change the biological properties of bacterial pathogens.

## Conflicts of interest

The authors declare that there are no conflicts of interest.

## Funding Information

This study is funded by the National Institute for Health Research (NIHR) Health Protection Research Unit in Genomics and Enabling Data at University of Warwick in partnership with the UK Health Security Agency (UKHSA), in collaboration with University of Cambridge and Oxford. Paolo Ribeca, Clare Barker and Marie Chattaway are based at UKHSA. Caitlin Collins is based at UKHSA and University of Cambridge. Xavier Didelot is based at University of Warwick. Matthew Bird is affiliated to the NIHR Health Protection Research Unit in Healthcare Associated Infections and Antimicrobial Resistance at University of Oxford in partnership with UKHSA; he is based at UKHSA. David Greig, Xavier Didelot, and Paolo Ribeca are affiliated to the NIHR Health Protection Research Unit in Gastrointestinal Infections at University of Liverpool in partnership with UKHSA, in collaboration with University of Warwick. David Greig is based at UKHSA. The views expressed are those of the author(s) and not necessarily those of the NIHR, the Department of Health and Social Care or the UK Health Security Agency. Paolo Ribeca acknowledges the Research/Scientific Computing teams at The James Hutton Institute and NIAB for providing computational resources and technical support for the UK’s Crop Diversity Bioinformatics HPC (BBSRC grant BB/S019669/1), use of which has contributed to the results reported within this article.

## Supplementary Figures

**Fig. S1.**
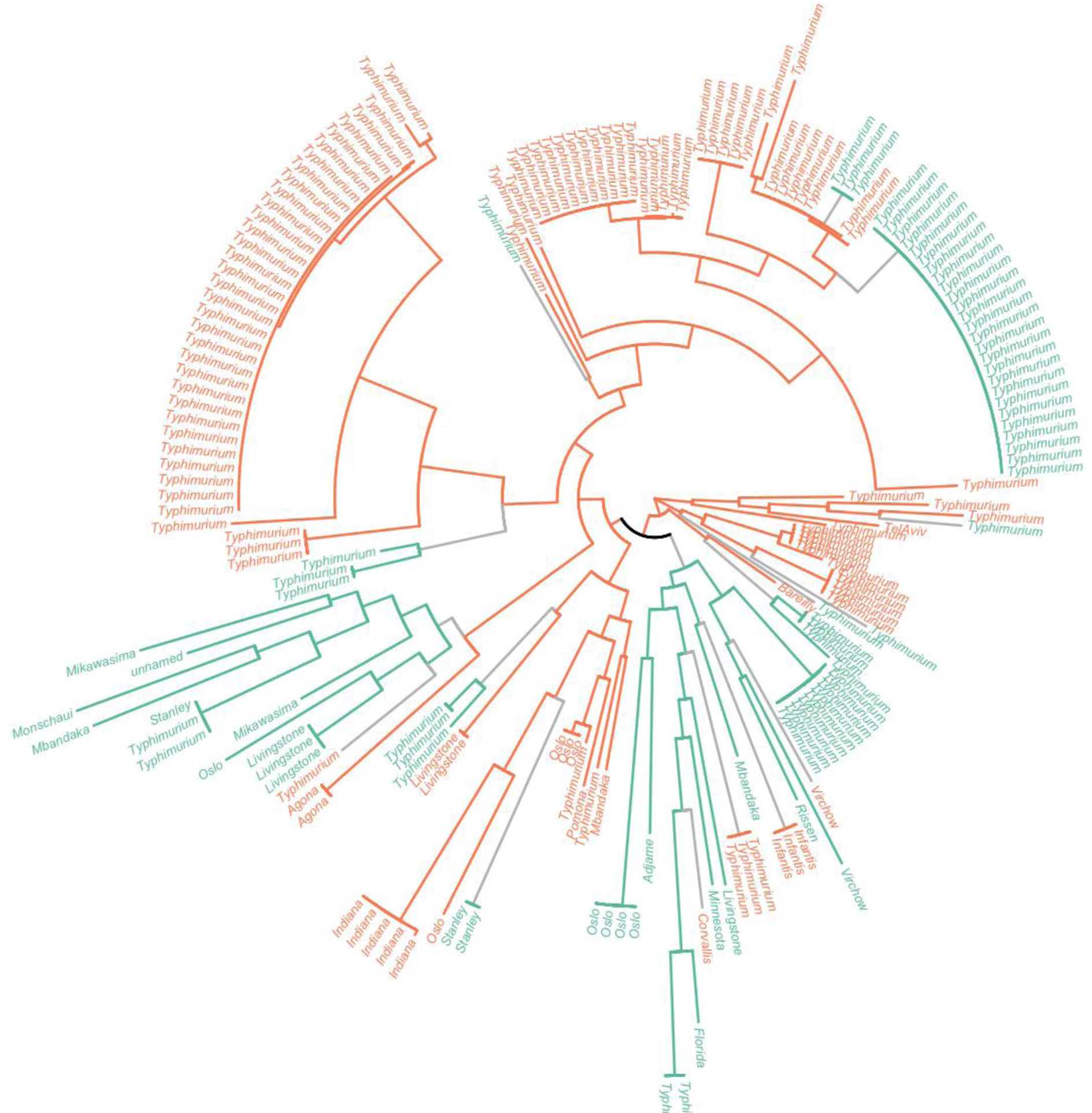
Phylogenetic tree based on the 51,260bp alignment of 54 core genes of the 193 *repL*-positive phage-plasmid contigs (Fig. 1) coloured by plasmid replication type, as observed at the tips and inferred via ancestral state reconstruction along the branches (IncY = green, p0111 = orange), showing 17 changes in replication type state (IncY>p0111 (5/17) or p0111>IncY (12/17) = grey).

**Fig. S2.**
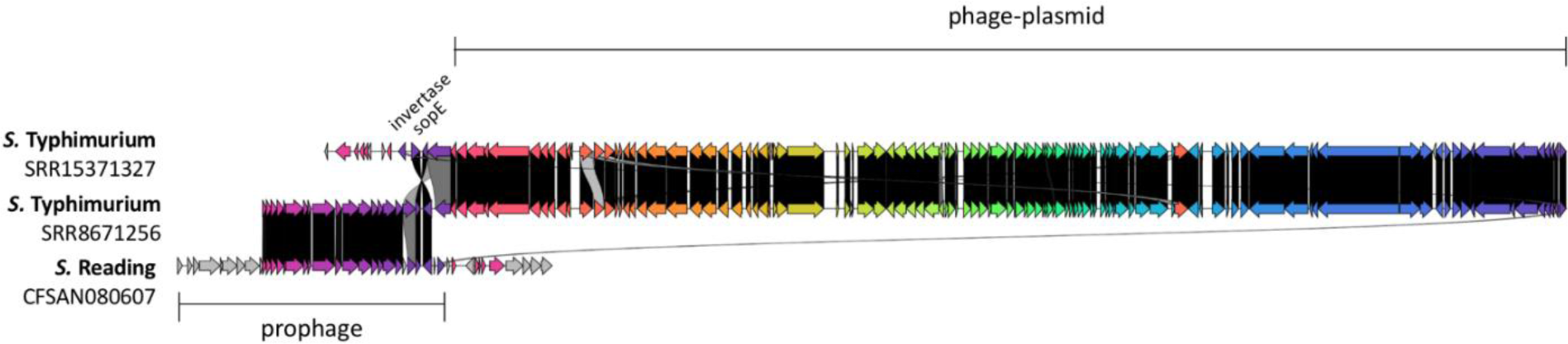
Phage-plasmids in *S.* Typhimurium putatively integrated in chromosome at site of prophage.

**Fig. S3.**
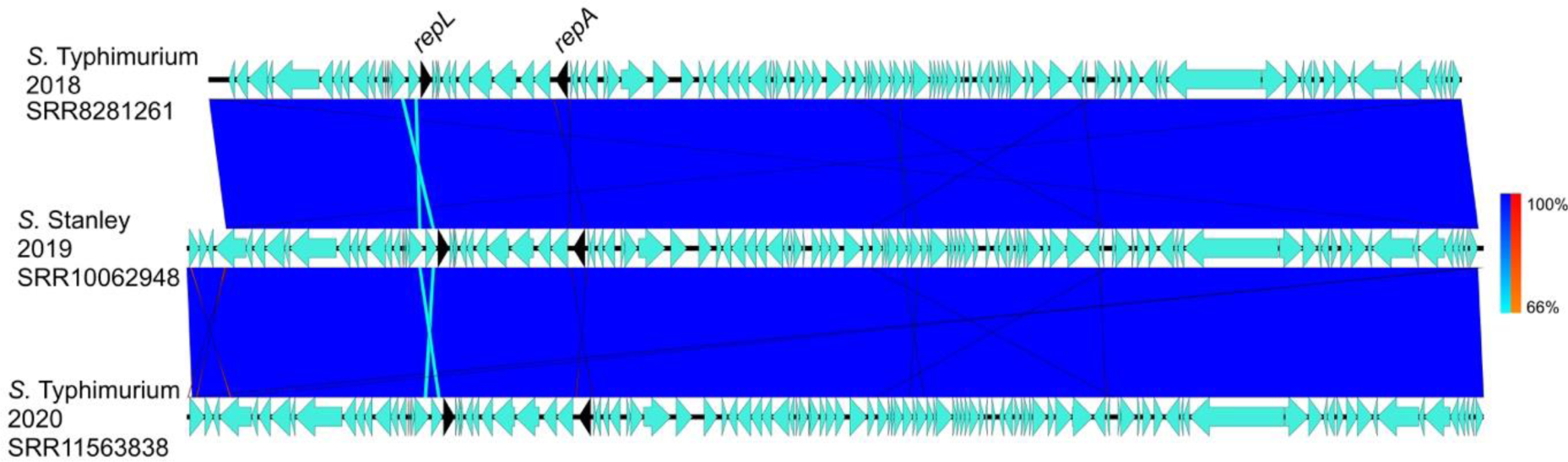
Highly similar phage-plasmid sequences in different serovars: *S.* Stanley from 2019, and *S.* Typhimurium from 2018 & 2020. Arrows indicate gene direction and scale bars indicate level of similarity for forward (blue) and reverse (red) sequences.

**Fig. S4.**
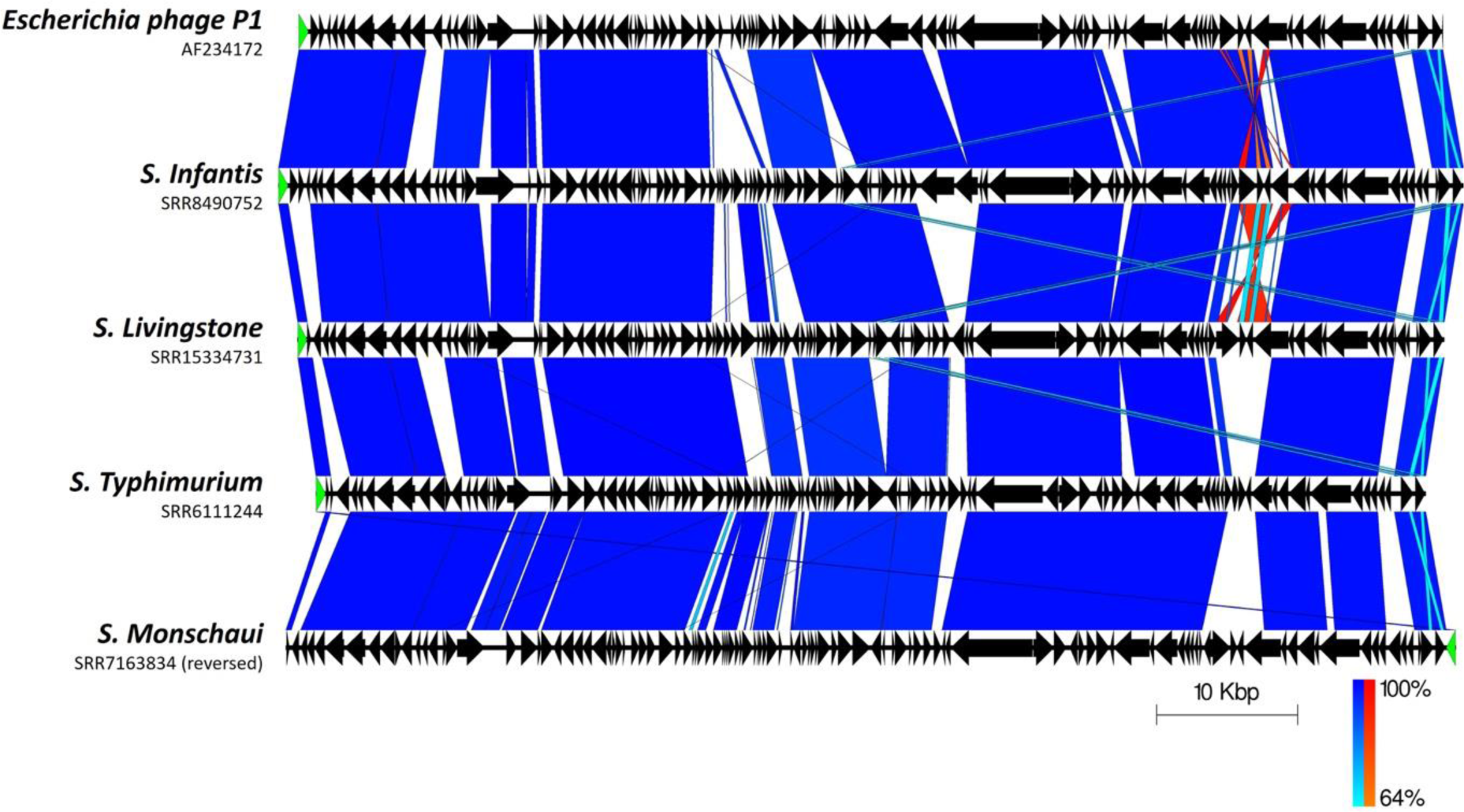
Variation in selected phage-plasmids from multiple serovars (*S.* Infantis, *S.* Livingstone*, S.* Monschaui and *S.* Typhimurium) compared to P1 bacteriophage. Arrows indicate gene direction and scale bars indicate level of similarity for forward (blue) and reverse (red) sequences. The *repL* gene is highlighted in green.

**Fig. S5.**
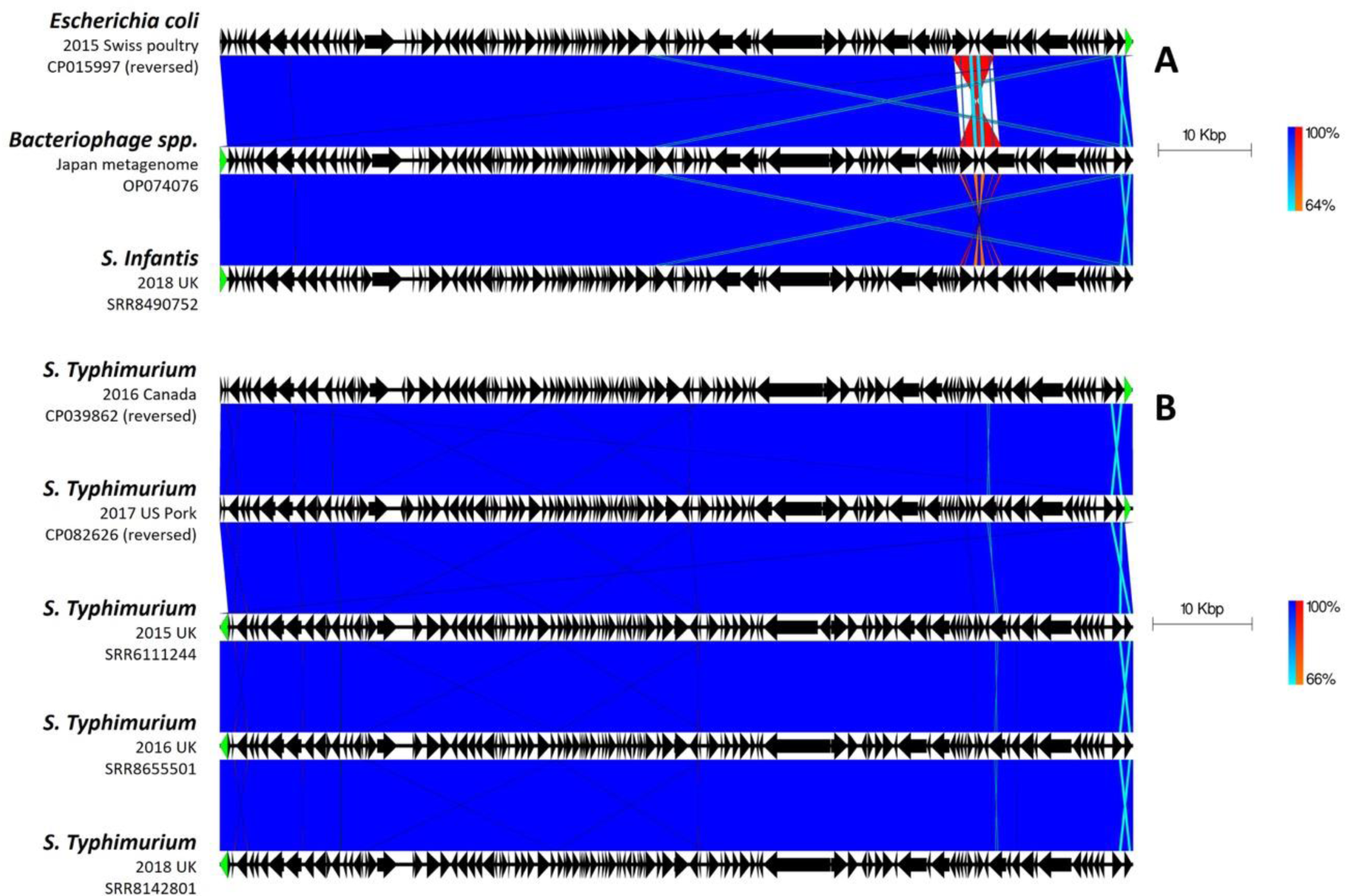
Highly similar phage-plasmid sequences from public databases showing conserved sequence across: **A)** different species; **B)** isolates of *S.* Typhimurium from different years and countries. Arrows indicate gene direction and scale bars indicate level of similarity for forward (blue) and reverse (red) sequences. The *repL* gene is highlighted in green.

**Fig. S6.**
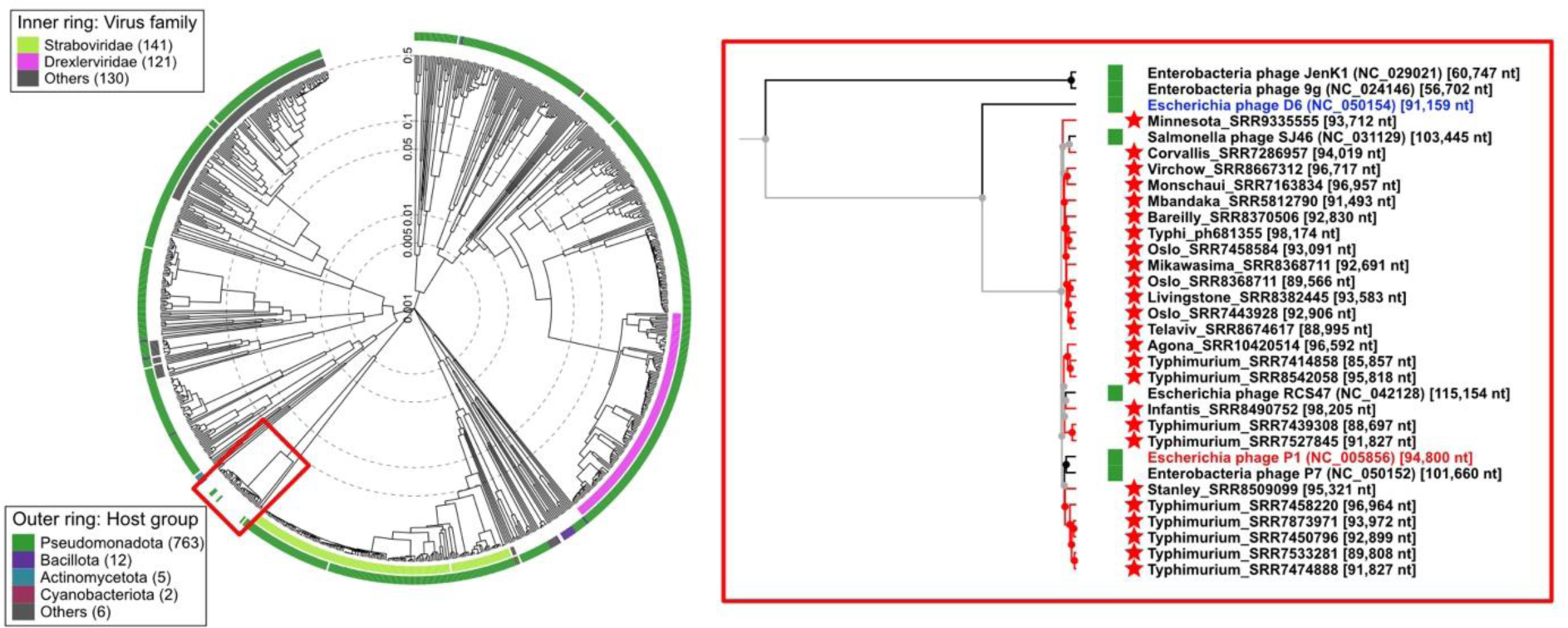
Detailed output from ViPTree server showing taxonomic context of P1-like phage-plasmids. Left: proteomic tree based on tBLASTx sequence similarity of 825 prokaryotic dsDNA viruses, with location of the P1-like clade highlighted in red; Right: zoomed-in view of highlighted clade, showing 25 representative phage-plasmids from this study and their nearest relatives such as P1 (red), D6 (blue), P7 and SJ46.

## Notes

### Competing Interest Statement

The authors have declared no competing interest.

### Summary of Updates

Text was revised, together with all figures, in order to improve clarity

## References

1. European Centre for Disease Prevention and Control., World Health Organization. Antimicrobial resistance surveillance in Europe 2022: 2020 data. [Internet]. LU: Publications Office; 2022 [cited 2023 Aug 2]. Available from: https://data.europa.eu/doi/10.2900/112339

2. European Food Safety Authority, European Centre for Disease Prevention and Control. The European Union Summary Report on Antimicrobial Resistance in zoonotic and indicator bacteria from humans, animals and food in 2019–2020. EFSA J [Internet]. 2022 Mar [cited 2023 Aug 2];20(3). Available from: https://data.europa.eu/doi/10.2903/j.efsa.2022.7209

3. Partridge SR, Kwong SM, Firth N, Jensen SO. Mobile Genetic Elements Associated with Antimicrobial Resistance. Clin Microbiol Rev. 2018 Aug;31(4):10.1128/cmr.00088-17.

4. Lindler LE, Plano GV, Burland V, Mayhew GF, Blattner FR. Complete DNA Sequence and Detailed Analysis of the Yersinia pestis KIM5 Plasmid Encoding Murine Toxin and Capsular Antigen. Infect Immun. 1998 Dec;66(12):5731–42.

5. Aviv G, Tsyba K, Steck N, Salmon-Divon M, Cornelius A, Rahav G, et al. A unique megaplasmid contributes to stress tolerance and pathogenicity of an emergent Salmonella enterica serovar Infantis strain. Environ Microbiol. 2014;16(4):977–94.

6. Brown-Jaque M, Calero-Cáceres W, Muniesa M. Transfer of antibiotic-resistance genes via phage-related mobile elements. Plasmid. 2015 May 1;79:1–7.

7. Balcazar JL. Bacteriophages as Vehicles for Antibiotic Resistance Genes in the Environment. PLOS Pathog. 2014 Jul 31;10(7):e1004219.

8. Brüssow H, Canchaya C, Hardt WD. Phages and the Evolution of Bacterial Pathogens: from Genomic Rearrangements to Lysogenic Conversion. Microbiol Mol Biol Rev. 2004 Sep;68(3):560–602.

9. Łobocka MB, Rose DJ, Plunkett G, Rusin M, Samojedny A, Lehnherr H, et al. Genome of Bacteriophage P1. J Bacteriol. 2004 Nov;186(21):7032–68.

10. Gilcrease EB, Casjens SR. The genome sequence of Escherichia coli tailed phage D6 and the diversity of Enterobacteriales circular plasmid prophages. Virology. 2018 Feb 1;515:203–14.

11. Pfeifer E, Moura de Sousa JA, Touchon M, Rocha EPC. Bacteria have numerous distinctive groups of phage–plasmids with conserved phage and variable plasmid gene repertoires. Nucleic Acids Res. 2021 Mar 18;49(5):2655–73.

12. Wang M, Jiang L, Wei J, Zhu H, Zhang J, Liu Z, et al. Similarities of P1-Like Phage Plasmids and Their Role in the Dissemination of blaCTX-M-55. Microbiol Spectr. 2022 Sep 7;10(5):e01410–22.

13. Pfeifer E, Bonnin RA, Rocha EPC. Phage-Plasmids Spread Antibiotic Resistance Genes through Infection and Lysogenic Conversion. mBio. 2022 Sep 26;13(5):e01851–22.

14. Chee MSJ, Serrano E, Chiang YN, Harling-Lee J, Man R, Bacigalupe R, et al. Dual pathogenicity island transfer by piggybacking lateral transduction. Cell. 2023 Aug 3;186(16):3414–3426.e16.

15. Billard-Pomares T, Fouteau S, Jacquet ME, Roche D, Barbe V, Castellanos M, et al. Characterization of a P1-Like Bacteriophage Carrying an SHV-2 Extended-Spectrum β-Lactamase from an Escherichia coli Strain. Antimicrob Agents Chemother. 2014 Nov;58(11):6550–7.

16. Bai L, Wang J, Hurley D, Yu Z, Wang L, Chen Q, et al. A novel disrupted mcr-1 gene and a lysogenized phage P1-like sequence detected from a large conjugative plasmid, cultured from a human atypical enteropathogenic Escherichia coli (aEPEC) recovered in China. J Antimicrob Chemother. 2017 May 1;72(5):1531–3.

17. Li R, Xie M, Lv J, Wai-Chi Chan E, Chen S. Complete genetic analysis of plasmids carrying mcr-1 and other resistance genes in an Escherichia coli isolate of animal origin. J Antimicrob Chemother. 2017 Mar 1;72(3):696–9.

18. Zhang C, Feng Y, Liu F, Jiang H, Qu Z, Lei M, et al. A Phage-Like IncY Plasmid Carrying the mcr-1 Gene in Escherichia coli from a Pig Farm in China. Antimicrob Agents Chemother. 2017 Feb 23;61(3):e02035–16.

19. Venturini C, Zingali T, Wyrsch ER, Bowring B, Iredell J, Partridge SR, et al. Diversity of P1 phage-like elements in multidrug resistant Escherichia coli. Sci Rep. 2019 Dec 11;9(1):18861.

20. Han H, Liu W, Cui X, Cheng X, Jiang X. Co-Existence of mcr-1 and blaNDM-5 in an Escherichia coli Strain Isolated from the Pharmaceutical Industry, WWTP. Infect Drug Resist. 2020 Apr 15;13:851–4.

21. Kalová A, Gelbíčová T, Overballe-Petersen S, Litrup E, Karpíšková R. Characterisation of Colistin-Resistant Enterobacterales and Acinetobacter Strains Carrying mcr Genes from Asian Aquaculture Products. Antibiotics. 2021 Jul;10(7):838.

22. Yao M, Zhu Q, Zou J, Shenkutie AM, Hu S, Qu J, et al. Genomic Characterization of a Uropathogenic Escherichia coli ST405 Isolate Harboring blaCTX-M-15-Encoding IncFIA-FIB Plasmid, blaCTX-M-24-Encoding IncI1 Plasmid, and Phage-Like Plasmid. Front Microbiol. 2022;13.

23. Zhou W, Liu L, Feng Y, Zong Z. A P7 Phage-Like Plasmid Carrying mcr-1 in an ST15 Klebsiella pneumoniae Clinical Isolate. Front Microbiol. 2018;9.

24. Yang L, Li W, Jiang GZ, Zhang WH, Ding HZ, Liu YH, et al. Characterization of a P1-like bacteriophage carrying CTX-M-27 in Salmonella spp. resistant to third generation cephalosporins isolated from pork in China. Sci Rep. 2017 Jan 18;7:40710.

25. Greig DR, Bird MT, Chattaway MA, Langridge GC, Waters EV, Ribeca P, et al. Characterization of a P1-bacteriophage-like plasmid (phage-plasmid) harbouring blaCTX-M-15 in Salmonella enterica serovar Typhi. Microb Genomics. 2022;8(12):000913.

26. Jiang L, Zhu H, Wei J, Jiang L, Li Y, Li R, et al. Enterobacteriaceae genome-wide analysis reveals roles for P1-like phage-plasmids in transmission of mcr-1, tetX4 and other antibiotic resistance genes. Genomics. 2023 Mar 1;115(2):110572.

27. Bolger AM, Lohse M, Usadel B. Trimmomatic: a flexible trimmer for Illumina sequence data. Bioinformatics. 2014 Aug 1;30(15):2114–20.

28. Achtman M, Wain J, Weill FX, Nair S, Zhou Z, Sangal V, et al. Multilocus Sequence Typing as a Replacement for Serotyping in Salmonella enterica. PLOS Pathog. 2012 Jun 21;8(6):e1002776.

29. Tewolde R, Dallman T, Schaefer U, Sheppard CL, Ashton P, Pichon B, et al. MOST: a modified MLST typing tool based on short read sequencing. PeerJ. 2016 Aug 17;4:e2308.

30. Dallman T, Ashton P, Schafer U, Jironkin A, Painset A, Shaaban S, et al. SnapperDB: a database solution for routine sequencing analysis of bacterial isolates. Bioinformatics. 2018 Sep 1;34(17):3028–9.

31. Marco-Sola S, Sammeth M, Guigó R, Ribeca P. The GEM mapper: fast, accurate and versatile alignment by filtration. Nat Methods. 2012 Dec;9(12):1185–8.

32. Li H, Handsaker B, Wysoker A, Fennell T, Ruan J, Homer N, et al. The Sequence Alignment/Map format and SAMtools. Bioinformatics. 2009 Aug 15;25(16):2078–9.

33. Danecek P, Bonfield JK, Liddle J, Marshall J, Ohan V, Pollard MO, et al. Twelve years of SAMtools and BCFtools. GigaScience. 2021 Feb 16;10(2):giab008.

34. Bonfield JK, Marshall J, Danecek P, Li H, Ohan V, Whitwham A, et al. HTSlib: C library for reading/writing high-throughput sequencing data. GigaScience. 2021 Feb 16;10(2):giab007.

35. Bankevich A, Nurk S, Antipov D, Gurevich AA, Dvorkin M, Kulikov AS, et al. SPAdes: A New Genome Assembly Algorithm and Its Applications to Single-Cell Sequencing. J Comput Biol. 2012 May;19(5):455–77.

36. Wick RR, Schultz MB, Zobel J, Holt KE. Bandage: interactive visualization of de novo genome assemblies. Bioinformatics. 2015 Oct 15;31(20):3350–2.

37. Camacho C, Coulouris G, Avagyan V, Ma N, Papadopoulos J, Bealer K, et al. BLAST+: architecture and applications. BMC Bioinformatics. 2009 Dec 15;10(1):421.

38. Bortolaia V, Kaas RS, Ruppe E, Roberts MC, Schwarz S, Cattoir V, et al. ResFinder 4.0 for predictions of phenotypes from genotypes. J Antimicrob Chemother. 2020 Dec 1;75(12):3491–500.

39. Tonkin-Hill G, MacAlasdair N, Ruis C, Weimann A, Horesh G, Lees JA, et al. Producing polished prokaryotic pangenomes with the Panaroo pipeline. Genome Biol. 2020 Jul 22;21(1):180.

40. Price MN, Dehal PS, Arkin AP. FastTree 2 – Approximately Maximum-Likelihood Trees for Large Alignments. PLOS ONE. 2010 Mar 10;5(3):e9490.

41. Argimón S, Abudahab K, Goater RJE, Fedosejev A, Bhai J, Glasner C, et al. Microreact: visualizing and sharing data for genomic epidemiology and phylogeography. Microb Genomics. 2016;2(11):e000093.

42. Schwengers O, Jelonek L, Dieckmann MA, Beyvers S, Blom J, Goesmann A. Bakta: rapid and standardized annotation of bacterial genomes via alignment-free sequence identification. Microb Genomics. 2021 Nov 5;7(11):000685.

43. Sullivan MJ, Petty NK, Beatson SA. Easyfig: a genome comparison visualizer. Bioinformatics. 2011 Apr 1;27(7):1009–10.

44. Gilchrist CLM, Chooi YH. clinker & clustermap.js: automatic generation of gene cluster comparison figures. Bioinformatics. 2021 Aug 25;37(16):2473–5.

45. Nishimura Y, Yoshida T, Kuronishi M, Uehara H, Ogata H, Goto S. ViPTree: the viral proteomic tree server. Bioinforma Oxf Engl. 2017 Aug 1;33(15):2379–80.

46. Joy JB, Liang RH, McCloskey RM, Nguyen T, Poon AFY. Ancestral Reconstruction. PLOS Comput Biol. 2016 Jul 12;12(7):e1004763.

47. R Core Team. R: A Language and Environment for Statistical Computing [Internet]. Vienna, Austria: R Foundation for Statistical Computing; 2012. Available from: https://www.R-project.org

48. Schliep KP. phangorn: phylogenetic analysis in R. Bioinformatics. 2011 Feb 15;27(4):592–3.

49. Collins C, Didelot X. A phylogenetic method to perform genome-wide association studies in microbes that accounts for population structure and recombination. PLOS Comput Biol. 2018 Feb 5;14(2):e1005958.

50. Mottawea W, Duceppe MO, Dupras AA, Usongo V, Jeukens J, Freschi L, et al. Salmonella enterica Prophage Sequence Profiles Reflect Genome Diversity and Can Be Used for High Discrimination Subtyping. Front Microbiol. 2018;9.

51. Kamal SM, Cimdins-Ahne A, Lee C, Li F, Martín-Rodríguez AJ, Seferbekova Z, et al. A recently isolated human commensal Escherichia coli ST10 clone member mediates enhanced thermotolerance and tetrathionate respiration on a P1 phage-derived IncY plasmid. Mol Microbiol. 2021;115(2):255–71.

52. Pfeifer E, Rocha EPC. Phage-plasmids promote recombination and emergence of phages and plasmids. Nat Commun. 2024 Feb 20;15(1):1545.

53. Bird MT, Greig DR, Nair S, Jenkins C, Godbole G, Gharbia SE. Use of Nanopore Sequencing to Characterise the Genomic Architecture of Mobile Genetic Elements Encoding blaCTX-M-15 in Escherichia coli Causing Travellers’ Diarrhoea. Front Microbiol [Internet]. 2022 [cited 2023 Sep 18];13. Available from: https://www.frontiersin.org/articles/10.3389/fmicb.2022.862234

54. Nair S, Chattaway M, Langridge GC, Gentle A, Day M, Ainsworth EV, et al. ESBL-producing strains isolated from imported cases of enteric fever in England and Wales reveal multiple chromosomal integrations of blaCTX-M-15 in XDR Salmonella Typhi. J Antimicrob Chemother. 2021 Jun 1;76(6):1459–66.

55. Zeng S, Luo J, Chen X, Huang L, Wu A, Zhuo C, et al. Molecular Epidemiology and Characteristics of CTX-M-55 Extended-Spectrum β-Lactamase-Producing Escherichia coli From Guangzhou, China. Front Microbiol [Internet]. 2021 [cited 2023 Sep 18];12. Available from: https://www.frontiersin.org/articles/10.3389/fmicb.2021.730012

56. Fernandes MR, Sellera FP, Cunha MPV, Lopes R, Cerdeira L, Lincopan N. Emergence of CTX-M-27-producing Escherichia coli of ST131 and clade C1-M27 in an impacted ecosystem with international maritime traffic in South America. J Antimicrob Chemother. 2020 Jun 1;75(6):1647–9.

57. Owen SV, Wenner N, Dulberger CL, Rodwell EV, Bowers-Barnard A, Quinones-Olvera N, et al. Prophages encode phage-defense systems with cognate self-immunity. Cell Host & Microbe. 2021 Nov 10;29(11):1620–1633.e8.

58. Georjon H, Bernheim A. The highly diverse antiphage defence systems of bacteria. Nat Rev Microbiol. 2023 Oct;21(10):686–700.

59. Fortier LC, Sekulovic O. Importance of prophages to evolution and virulence of bacterial pathogens. Virulence. 2013 Jul 1;4(5):354–65.

60. Jaslow SL, Gibbs KD, Fricke WF, Wang L, Pittman KJ, Mammel MK, et al. Salmonella Activation of STAT3 Signaling by SarA Effector Promotes Intracellular Replication and Production of IL-10. Cell Reports. 2018 Jun 19;23(12):3525–36.

61. Schroven K, Aertsen A, Lavigne R. Bacteriophages as drivers of bacterial virulence and their potential for biotechnological exploitation. FEMS Microbiology Reviews. 2021 Jan 8;45(1):fuaa041.

62. Andrews K, Landeryou T, Sicheritz-Pontén T, Nale JY. Diverse Prophage Elements of Salmonella enterica Serovars Show Potential Roles in Bacterial Pathogenicity. Cells. 2024 Jan;13(6):514.

63. Bondy-Denomy J, Davidson AR. When a virus is not a parasite: the beneficial effects of prophages on bacterial fitness. J Microbiol. 2014 Mar 1;52(3):235–42.

64. Fong K, Lu YT, Brenner T, Falardeau J, Wang S. Prophage Diversity Across Salmonella and Verotoxin-Producing Escherichia coli in Agricultural Niches of British Columbia, Canada. Front Microbiol. 2022 Jul 22;13.

65. Figueroa-Bossi N, Uzzau S, Maloriol D, Bossi L. Variable assortment of prophages provides a transferable repertoire of pathogenic determinants in Salmonella. Mol Microbiol. 2001;39(2):260–72.

66. Octavia S, Sara J, Lan R. Characterization of a large novel phage-like plasmid in Salmonella enterica serovar Typhimurium. FEMS Microbiol Lett. 2015 Apr 1;362(8):fnv044.

67. Colavecchio A, Jeukens J, Freschi L, Edmond Rheault JG, Kukavica-Ibrulj I, Levesque R, et al. AnCo3, a New Member of the Emerging Family of Phage-Like Plasmids. Genome Announc. 2017 May 11;5(19):10.1128/genomea.00110-17.

68. Colavecchio A, Jeukens J, Freschi L, Edmond Rheault JG, Kukavica-Ibrulj I, Levesque RC, et al. Complete Genome Sequences of Two Phage-Like Plasmids Carrying the CTX-M-15 Extended-Spectrum β-Lactamase Gene. Genome Announc. 2017 May 11;5(19):10.1128/genomea.00102-17.

69. Katz K, Shutov O, Lapoint R, Kimelman M, Brister JR, O’Sullivan C. The Sequence Read Archive: a decade more of explosive growth. Nucleic Acids Res. 2022 Jan 7;50(D1):D387–90.

70. Zhou Z, Alikhan NF, Mohamed K, Fan Y, Group the AS, Achtman M, et al. The EnteroBase user’s guide, with case studies on Salmonella transmissions, Yersinia pestis phylogeny, and Escherichia core genomic diversity. Genome Res. 2020 Jan 1;30(1):138–52.

